# Efficacy of aerial forward-looking infrared surveys for detecting polar bear maternal dens

**DOI:** 10.1101/763144

**Authors:** Tom S. Smith, Steven C. Amstrup, John Kirschhoffer, Geoffrey York

## Abstract

Denned polar bears are invisible under the snow, therefore winter-time petroleum exploration and development activities in northern Alaska have potential to disturb maternal polar bears and their cubs. Previous research determined forward looking infrared (FLIR) imagery could detect many polar bear maternal dens under the snow, but also identified limitations of FLIR imagery. We evaluated the efficacy of FLIR-surveys conducted by oil-field operators from 2004-2016. Aerial FLIR surveys detected 15 of 33 (45%) and missed 18 (55%) of the dens known to be within surveyed areas. While greater adherence to previously recommended protocols may improve FLIR detection rates, the physical characteristics of polar bear maternal dens, increasing frequencies of weather unsuitable for FLIR detections—caused by global warming, and competing “hot spots” are likely to prevent FLIR surveys from detecting maternal dens reliably enough to afford protections consonant with increasing global threats to polar bear welfare.

## Introduction

Polar bears construct snow dens in which they give birth to and nurture altricial young. In arctic Alaska, dens in drifted snow are excavated from mid-October through early December. Cubs are born in midwinter (Amstrup 2003), and family groups abandon dens by mid-April (Amstrup 1993 and Smith et al. 2007).

The geographic scope of petroleum exploration and development in the Alaskan Beaufort Sea coastal region has been expanding, and is now even proposed for the Coastal Plain of the Arctic National Wildlife Refuge (https://eplanning.blm.gov/epl-front-office/eplanning/planAndProjectSite.do?methodName=dispatchToPatternPage&currentPageId=152110) which is designated critical polar bear denning habitat. Simultaneously, the proportion of maternal polar bears choosing to den on land has been increasing (Amstrup and Gardner 1994, Fischbach et al. 2007). The likelihood that maternal dens could be disturbed therefore, can be expected to increase. Because polar bear cubs are born very altricial and cannot leave the shelter of the den until approximately 3 months of age (Amstrup 2003), disruption of denning can have negative consequences for cubs and maternal females (Amstrup 1993, Linnell et al. 2000). Additionally, industry-related den disturbance can have significant economic consequences, including: rerouting roads, delays in exploration and production, fines, and other penalties.

Maternal dens usually remain unopened through winter and are essentially invisible under the snow. Previous research determined forward looking infrared (FLIR) imagery could detect the temperature differential between snow over some polar bear maternal dens and the snow where dens were absent. That research, however, also identified limitations of FLIR imagery, and recommended “best practices” protocols to maximize detection abilities (Amstrup et al. 2004, York et al. 2004, Robinson et al. 2014). To detect and hence avoid disturbances to maternal dens, oil companies operating in southern Beaufort Sea coastal areas of northern Alaska began using FLIR in 2004 to locate polar bear dens within oil-field operating areas so they can be avoided during ice road construction and other exploration and production activities.

The purpose of this study was to evaluate the efficacy of industry-operated aerial FLIR surveys (hereafter referred to as “industry AFS”) and make recommendations for future den detection and avoidance efforts.

### Study Area

The study area included northern Alaska’s Beaufort Sea coastal areas (commonly called the North Slope), extending 133 km west and 91 km east of Prudhoe Bay (70°20’ N, 148°24’ W, Figure 1). The Prudhoe Bay region has a semi-arid-tundra climate. The mean annual temperature at Deadhorse, an unincorporated community providing airport services, weather observations and an unofficial hub for Prudhoe Bay operations, is −11 °C (12 °F). The warmest month, July, has a daily average temperature of 8.3 °C (47 °F), the coldest month February at −28 °C (−18 °F; Deadhorse 2019).

**Fig. 1.**
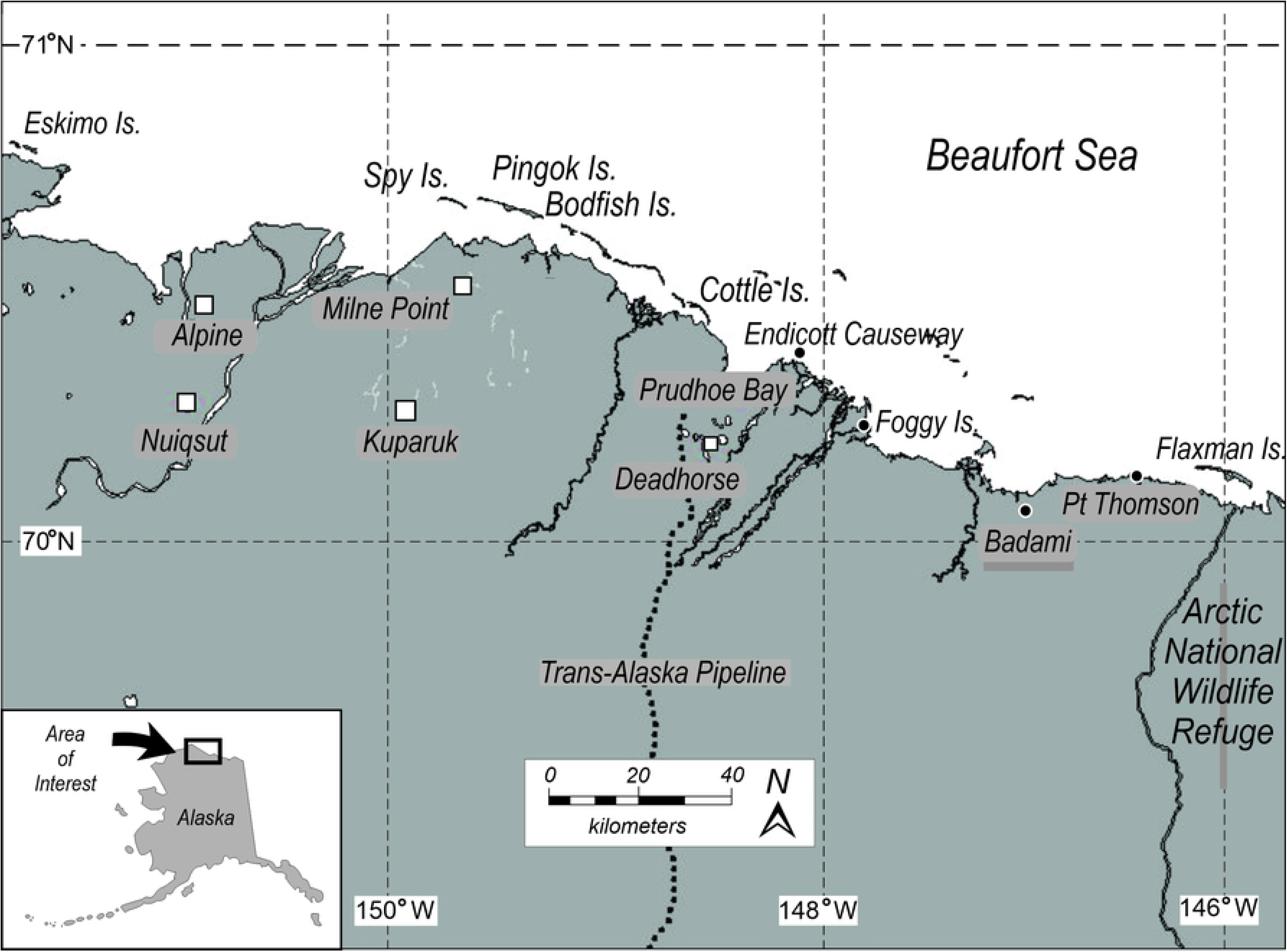
Study area where FLIR aerial surveys and ground truthing of polar bear den detection surveys were conducted in northern Alaska, 2004-2016.

Alaska’s North Slope lacks the steep topography associated with other denning areas such as Wrangel and Herald Islands, Russia (Uspenski and Kistchinski, 1972, Ovsyanikov, 1998), and Svalbard, Norway (Larsen, 1985). The predominantly flat topography of coastal arctic Alaska means suitable denning habitat is restricted to riverbanks, coastal bluffs, barrier islands and other areas where relief is sufficient to catch drifting snow (Amstrup 1993, Amstrup and Gardner 1994, Durner et al. 2001, Durner et al. 2003, Durner et al. 2006). Amstrup (2003) reported over 80% of dens, located by radio telemetry along Alaska’s north slope, were within 10km of shore, but a small number have been located as far inland as 50 km (Durner et al., 2003).

## Methods

We assessed the efficacy of industry AFS for polar bear den site detection by comparing industry AFS results, for the period 2004-2016, with ground-truth data we collected during research we conducted on emergence behaviors of denning polar bears between 2002 and 2016 in the same area (Smith et al. 2007, Smith et al. 2013). Our on the ground documentation of actual den sites provided ground truth data against which industry AFS results were compared for this report.

Industry AFS were conducted with the Star Safire (models II, III, and HD 380 FLIR camera units (www.flir.com). FLIR units employed were gimbal-mounted (single axis rotational support) under a de Havilland DHC-6 Twin Otter for all surveys referenced here. This mounting system allowed the FLIR imager to be directed independent of altitude and in any direction below the horizontal plane of the aircraft. The Safire, operates in the 8 to 14 micron wavelength range, and under ideal circumstances can detect temperature differences as small as 0.1°C (FLIR Systems). Here we refer to thermal signatures detected by industry AFS as “hot spots.” When a hot spot was detected, observers changed FLIR camera view angle, aircraft altitude and position in an attempt to determine whether it was a den or some other source of heat. Numerous potential targets in this environment (e.g. cracks in sea ice, exposed soil, large rocks, or some manmade objects like abandoned 55-gallon steel drums) collect and reradiate heat differently than snow covered ground, and can emit thermal signals similar to those from dens (Figure 2). Hotspots with the “right” signature, detected during surveys, were marked as putative maternal dens. FLIR generated video, including observer audio comments about detected hotspots, was recorded for each flight and archived. Surveyors recorded weather conditions, reported by the flight service station in Deadhorse, during each survey. To supplement weather data recorded during industry AFS, we collected data from the closest ground-based weather stations to each survey flight. These data included ambient temperature, wind speed, relative humidity, and dew point. Dew point, the temperature to which a given parcel of air must be cooled for saturation to occur, incorporates the effect of pressure and temperature on relative humidity.

**Fig. 2.**
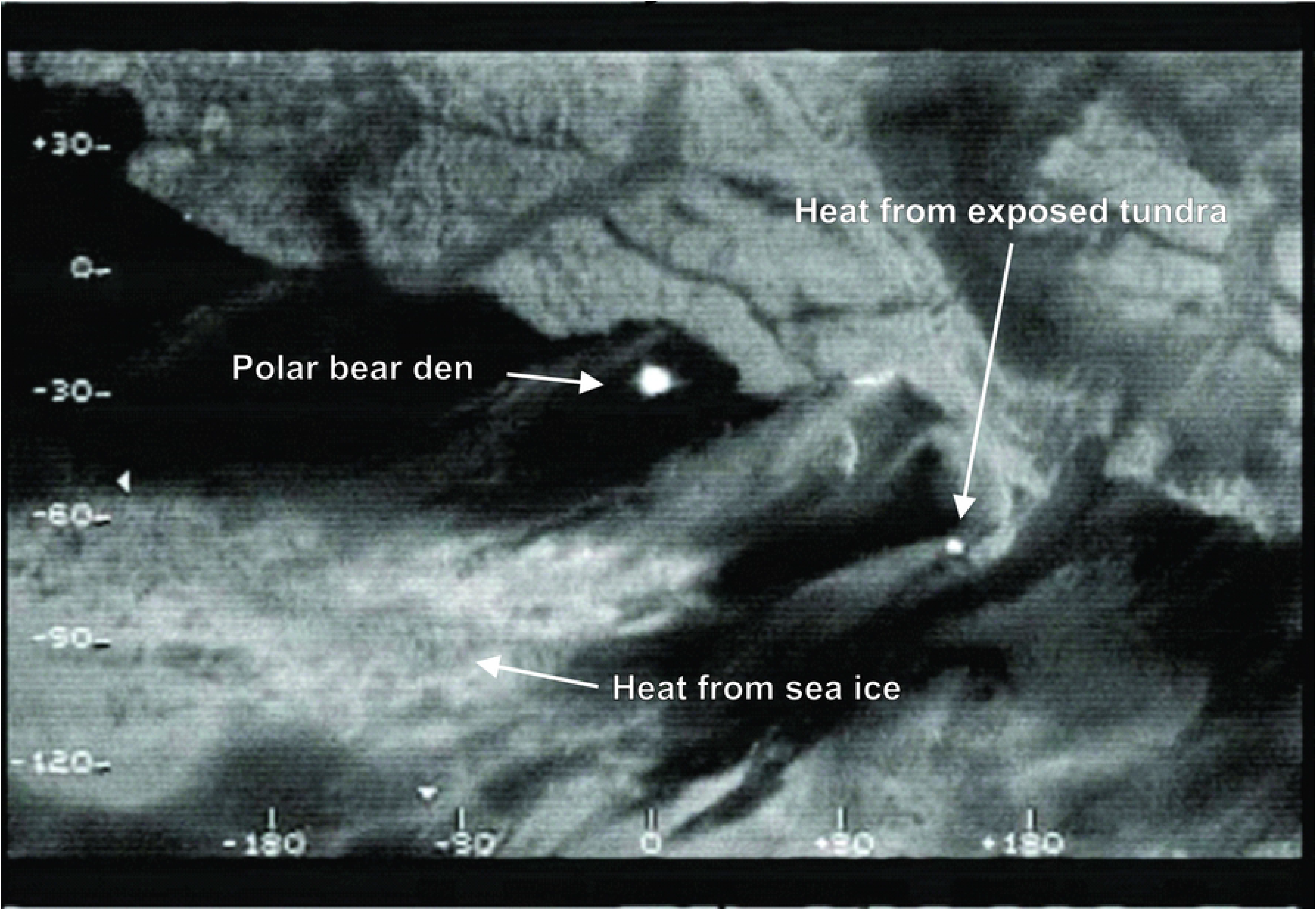
Forward-looking infrared image of two polar bear dens in the snow bank on the south shore of an Alaskan coastal island. Also note hotspots created by exposed tundra and warmth radiating up from sea ice with a thin covering of snow.

Personal involved in industry AFS provided us with complete survey reports for the years 2008-2016. Those reports provide dates, times, prevailing weather conditions, number of putative dens recorded, and areas in which no dens were observed. Prior to 2008, we received only putative den locations and observations of other polar bear signs.

Along with putative den locations from industry AFS, we used a hand-held FLIR imager (ThermaCAM P65 HS, FLIR Systems) with a 72mm infrared telephoto lens to identify potential den sites by visiting historically high-use denning areas by snowmachine and scanning snowbanks with the hand-held unit. Additionally, den locations of radio-collared polar bears were provided by the U.S. Geological Survey (USGS) and by the US Fish and Wildlife Service (FWS) that had been confirmed by visual observation or the use of Karelian Bear Dogs. We monitored putative den locations provided by industry AFS, to ascertain which ones were correctly identified as maternal dens. In addition to positive FLIR identifications of den sites, we tabulated the number of dens known to be within surveyed areas that were missed by industry AFS (false negatives), and we evaluated the frequency of hotspots that were incorrectly identified as dens but proved otherwise (false positives). After bears left their dens in spring, we visited each confirmed den site and recorded the depth of snow overlying the main chamber, and other site characteristics (consistent with Durner et al. 2003).

## Results

Once den breakout began in the study area (mean date = 16 March, range = 1 March to 4 April; Figure 3), den locations could be verified visually because piles of excavated snow and tracks and other signs of bear activity surrounding den openings were visible on the snow surface. Consequently, we believe our thorough ground assessment of the survey area provided an accurate evaluation of industry AFS results.

**Fig. 3.**
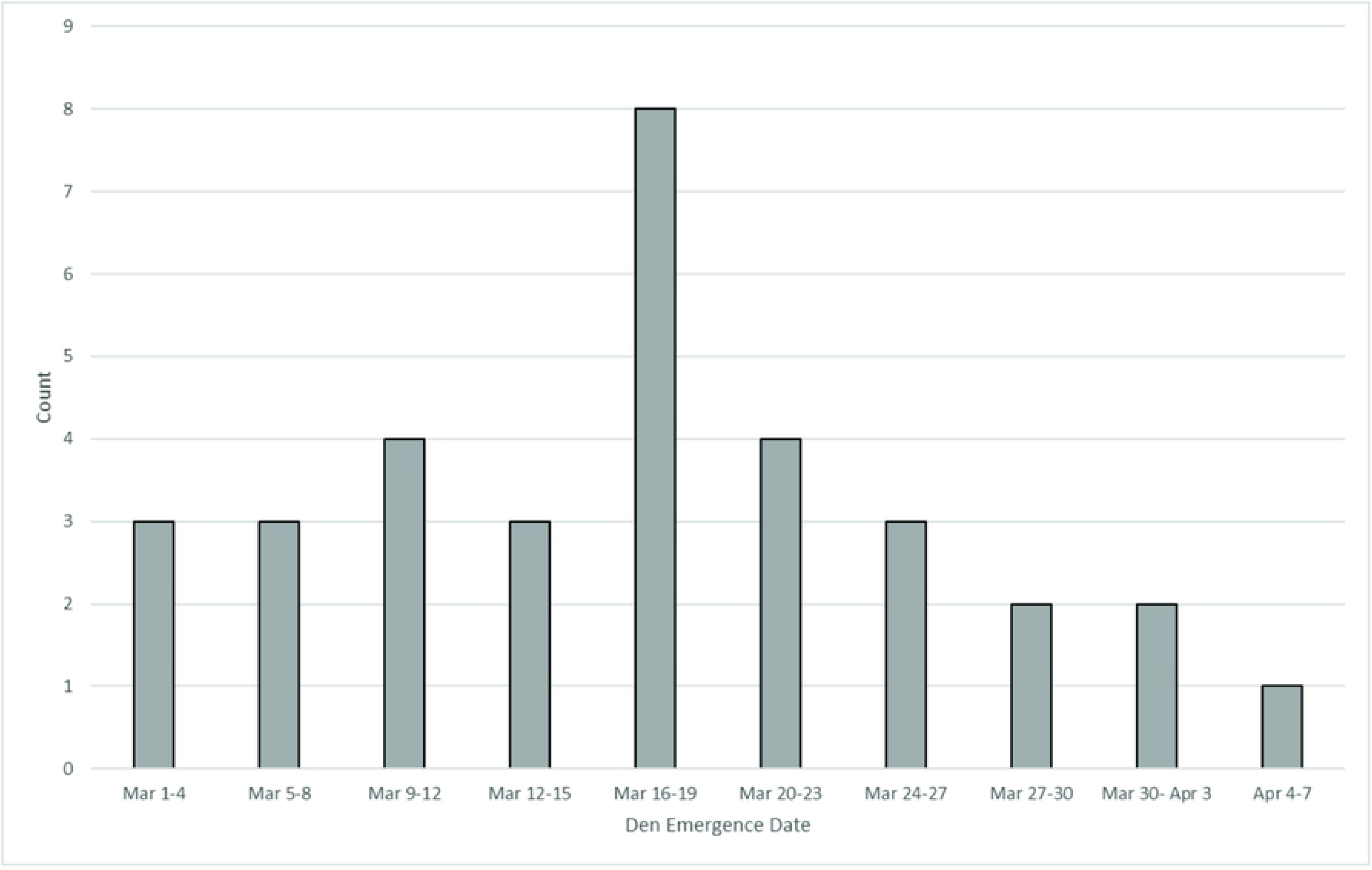
Emergence dates for 32 polar bear dens within the study area from 2002-2014. Emergence time for one den discovered during this study was not accurately known.

Between 2004 and 2016, we identified 33 maternal polar bear dens within areas also surveyed by industry. Of those 33 dens, 15 (45%) were accurately detected by industry AFS (Table 1). Industry AFS operators also identified 19 putative polar bear dens that our field work proved to be thermal signatures generated by other sources. Because aerial inspection of these “false positives,” during industry AFS, proved insufficient to differentiate them from real dens, monitoring and activity restrictions required for maternal dens also were required for these locations. Prevailing weather at the time of each AFS is presented in Table 2. No industry AFS den surveys were conducted when the sun was above the horizon.

**Table 1.**
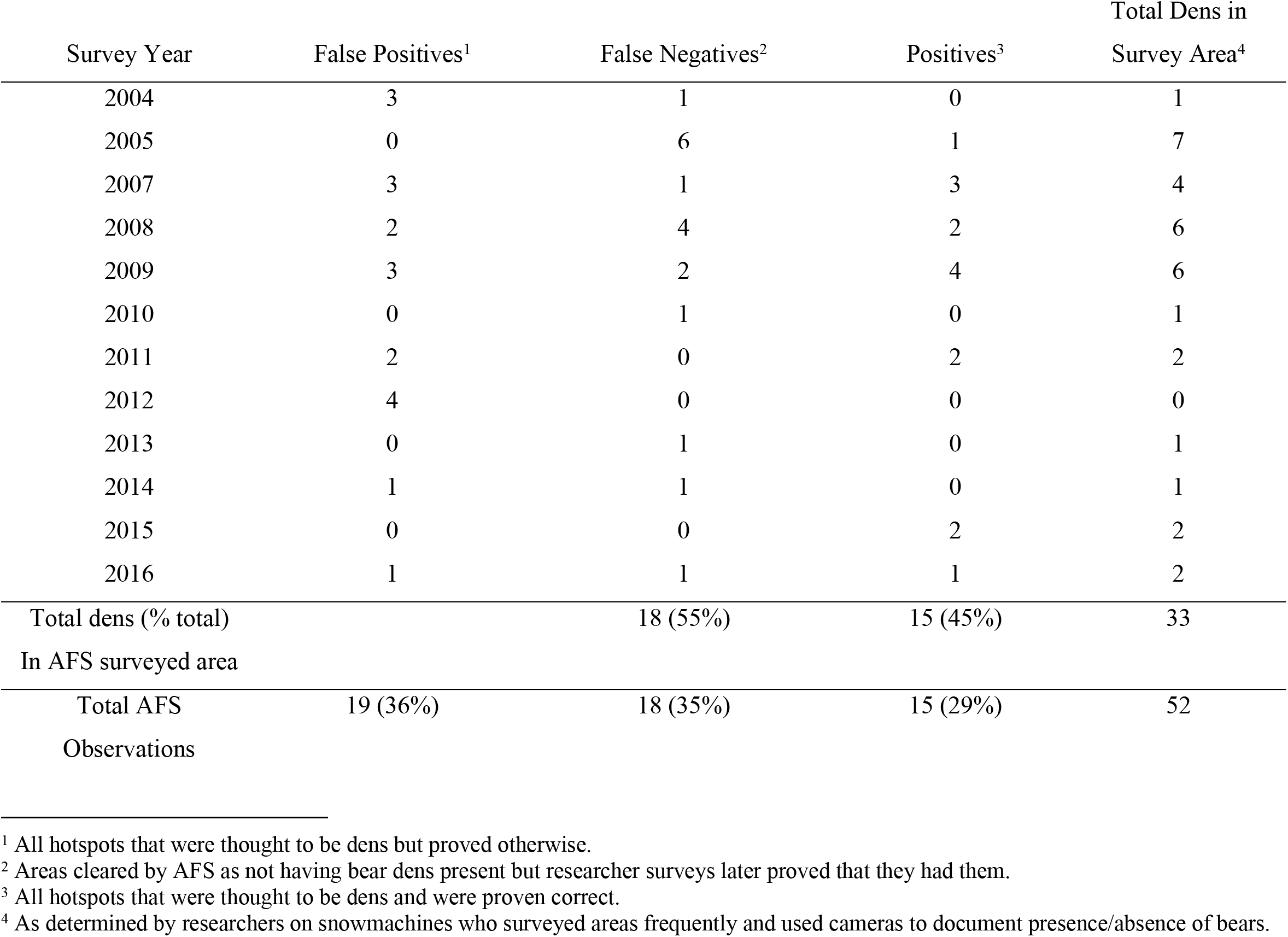
Summary of observations collected during aerial FLIR surveys to detect polar bear dens in northern Alaska, 2004 to 2016.

**Table 2.** Prevailing weather conditions during FLIR aerial survey flights for polar bear den detection, northern Alaska, 2007—2016. We were not provided weather data for surveys conducted prior to 2007.

| Survey Date | Temperature<br>(°C) | Dew point/temp spread<br>(°C) | Wind Speed<br>(km/h) | Misidentified<br>Den Sites |
| --- | --- | --- | --- | --- |
| 4 Dec 2007 | -15 | 5 | 9.7 | 3 |
| 29 Jan 2008 | -24 | 19 | 29.0 | 3 |
| 8 Dec 2008 | -22 | no data | 33.8 | 3 |
| 7 Dec 2009 | -8 | 12 | 0.2 | 5 |
| 17 Dec 2011 | -24 | 3 | 32.2 | 2 |
| 6 Jan 2012 | -27 | 3 | 16.1 | 4 |
| 14 Dec 2012 | -38 | no data | 9.7 | 1 |
| 9 Dec 2013 | -15 | 2 | 48.3 | 3 |
| 3 Dec 2014 | -11 | 7 | 3.2 | 1 |
| 4 Dec 2014 | -6 | 7 | 6.4 | 1 |
| 5 Dec 2014 | 3 | 5 | 16.1 | 1 |
| 6 Dec 2014 | -8 | 5 | 25.7 | 2 |
| 4 Dec 2015 | -28 | 3 | 16.1 | 2 |
| | $\bar{x} = -17$ | $\bar{x} = 6.5$ | $\bar{x} = 19.0$ | Total = 31 |

## Discussion

During 13 years of industry AFS only 15 of 33 (45% detection rate) polar bear dens known to be within the areas surveyed were detected. One possible explanation for failure of industry AFS to detect dens is that some female bears could have entered dens late in the season—after the AFS was conducted. However, Amstrup and Gardner (1994) reported November 11 was the mean den entrance date for polar bears denning successfully on land. Rode et al. (2018), also reported a November 11 mean den entrance date for the polar bears in northern Alaska with a ± SD of 18.5 days. While it is possible for bears to have entered dens after survey dates presented in Table 2, nearly 95% of all bears were denned before 4 December (USGS 2018)—the earliest recorded industry AFS effort. Hence ambient conditions and other limitations of FLIR are more likely drivers of the low detection rate of industry AFS.

In order to test den detection effectiveness of AFS, Amstrup et al. (2004) flew multiple FLIR surveys over 23 denning bears for which exact locations were known by radio-telemetry. Only 7 of the 23 dens (30%) were detected on every flight, and 4 dens (17%) were never detected. Weather conditions (e.g., wind, precipitation, temperature-dew point spread) and conducting surveys in daylight or to soon after snow fall or drifting caused by high wind, were identified as principal reasons for detection failure. Robinson et al. (2014) recommended against conducting surveys during daylight hours and Amstrup et al. (2004) concluded the probability of detecting dens in sunlight was essentially zero. Robinson et al. (2014) also reported that when wind was > 10 km/h den detection with FLIR was unlikely. Of the industry AFS listed in Table 2, 42% (5 of 12) were conducted with winds < 10 km/h. However, of those five surveys, two were conducted with prevailing winds very close to the detectability cutoff (9.7 km/h). Hence, only 25% (*n* = 3) of industry AFS, for which we have data on ambient conditions, were performed under wind conditions that were conducive to den detection with FLIR.

Although ambient conditions at the time of surveys appear to explain much of the detection failure rate; the 4 dens Amstrup et al. (2004) never detected were visited 6 times, including FLIR surveys conducted under optimal weather conditions. Variable snow depth over maternal dens may explain detection failures occurring even when survey conditions seem appropriate. Robinson et al. (2014) reported that even under ideal ambient conditions, hand-held FLIR was unable to detect a thermal signature emanating from artificial test dens if roof thickness was 90 cm or greater. While the actual roof thickness that precludes detection by industry AFS is uncertain, it is clear there is a snow depth threshold that prevents sufficient heat generated by denned bears from reaching the snow surface. Durner et al. (2003) reported the mean den roof thicknesses for 22 polar bear dens in northern Alaska was 72 ± 87 cm, and ranged from as little as 10cm to as much as 400 cm. Snow depth over many dens, therefore, is likely near or above FLIR detection capabilities, regardless of weather—corroborating the conclusion of Amstrup et al. (2004) that some dens will never be detected with FLIR.

Mid-winter den abandonment may account for some putative den locations that we found not to be actual dens. More hotspots not associated with den emergence (36% of all putative den sites, Table 1), were reported by industry AFS than the actual number of dens detected. This far exceeds documented mid-winter den abandonments (mean = 12 % across all years, USGS 2018). York et al. (2004) advised that hotspots of interest should be revisited on subsequent days under good environmental conditions to confirm whether they are truly dens.. The rate of these “false positive” signals far exceeds anything that could be expected because of mid-winter den abandonment, suggesting industry AFS did not adhere to protocols known to minimize false positives.

## Conclusions

The U.S. Fish and Wildlife Service Conservation Management Plan (U.S. Fish and Wildlife 2016) recognizes the need for “on the ground” protections to assure as many polar bears as possible persist until sea ice is stabilized. The catastrophic decline (~40% between 2000 and 2010) in the Southern Beaufort Sea polar bear population was driven by reduced survival, particularly of cubs (Bromaghin et al. 2015). This makes it clear that maximizing cub survival potential is essential to maximizing opportunity for polar bears in this region to persist. A critical step toward maximizing cub survival is protecting maternal dens. Where denning activity overlaps with intensive human activities like oil and gas development, protection of maternal dens begins with knowing where they are. By that measure, industry AFS has fallen short of the protections the imperiled southern Beaufort Sea population of polar bears requires.

The poor den detection rate of industry AFS is likely due to a combination of factors, including weather-related variables (e.g., wind, temperature-dew point spread, precipitation, etc.), time of day, and den roof thickness (Table 2). The latest generation of FLIR imagers, while more advanced and sensitive than those used in the earlier surveys, will still struggle with the fundamental physics of detecting subsurface heating associated with dens in the presence of strong winds, direct solar radiation, falling or blowing snow, or too deep snow overlaying the den (R. Overstreet, FLIR Sales Engineer, personal communications, 21 February 2019). Available data make it clear that FLIR will never assure that all maternal dens can be located and hence protected. In addition, as the Arctic has warmed airborne moisture has increased and is expected to continue to do so (Serreze et al. 2012). Because airborne moisture significantly limits FLIR detection capabilities, finding optimal operational windows for FLIR surveys, is likely to become increasingly difficult, and probabilities of den detection by FLIR are likely only to decline. Because global temperature rise ultimately will impact all polar bears, and because FLIR is unlikely to offer necessary protections, developing other technologies to protect maternal denning bears is of utmost importance.

To maximize detection rate in the near term, industry AFS should consistently follow recommended survey protocols developed during previous research (Amstrup et al. 2004, York et al. 2004, Robinson et al. 2014). Surveys should be done 1) with the sun below the horizon, 2) with a temperature-dew point spread > 2.8 °C, 3) with winds < 10 km/h, 4) without precipitation, 5) not immediately after a strong wind event, 6) with helicopters rather than fixed-wing aircraft, 7) and with multiple flights to eliminate hotspots that are not dens. Also, to the maximum extent possible, industry should continue to conduct AFS in early winter (i.e., early December), when snow accumulation over dens is likely to be at winter minimums. While best FLIR practices are applied in the near term, we encourage the exploration of other technologies that could increase our ability to detect polar bear dens. Synthetic aperture radar (SAR) has shown promise for detecting dens, and is not vulnerable to the weather and daylight constraints that limit FLIR application. Significant testing, however, is required to properly evaluate the promise of SAR. Regardless of whether SAR surveys are the answer, the current and future threats to polar bears mandate development of new den location technologies with higher detection rates than possible with FLIR.

## Acknowledgements

We would like to thank anonymous reviewers for their thoughtful comments regarding this manuscript. We would also like to thank the following persons for assisting data collection: S. Partridge, J. Wilder, T. D. DeBruyn, J. Wilder, T. Evans, S. Schliebe, C. Perham, R. Shideler, B. Jessop, J. Olson, R. Robinson, W. G. Larson and J. Whiting. Additionally, we thank W. J. Streever, D. Sanzone, and C. Pohl of British Petroleum Exploration (BPX) for their support and assistance, without which this work would not have been possible. Also important was R. Murray and N. Hermon of Alaska Clean Seas, and S. Gogosha and D. Heebner of HilCorp Energy Company who provided invaluable assistance at the Milne Point facility. We extend special thanks to D. Herron and J. Thompson of the BP Thermographics Division for providing access to the FLIR cameras. Polar Bears International (PBI) has played a critical role in supporting this work. Funding for this work was provided, in part by the U.S. Fish and Wildlife Service, the U.S. Geological Survey, BPX, PBI and BYU.

## References

1. Amstrup, S.C. Human disturbances of denning polar bears in Alaska. Arctic 1993; 46: 246–250.

2. Amstrup, S.C., and Gardner, C. Polar bear maternity denning in the Beaufort Sea. J. Wildl. Manage. 1994; 58: 1–10.

3. Amstrup, S. C. The polar bear, *Ursus maritimus*. Chapter 27 in Wild Mammals of North America: biology, management, and conservation. Edited by G.A. Feldhamer, B.C. Thompson, and J.A. Chapman. John Hopkins Univ. Press, Baltimore. 2003; pp. 587–610.

4. Amstrup, S. C., G. York, T. L. McDonald, R. Nielson, and K. Simac. Detecting denning polar bears with forward looking infra-red (FLIR) imagery. BioSci. 2004; 54: 337–344.

5. Bromaghin, J.F., T.L. Mcdonald, I. Stirling, A. E. Derocher, E.S. Richardson, E.V. Regehr, D. C. Douglas, G.M. Durner, T. Atwood, and S.C. Amstrup. Polar bear population dynamics in the Southern Beaufort Sea during a period of sea ice decline. Eco. Appl. 2015; 25: 634–651.

6. Deadhorse, Alaska. Wikipedia. Wikimedia Foundation. Available from: https://en.wikipedia.org/wiki/Deadhorse,_Alaska.

7. Durner, G. M., S. C. Amstrup, and K. J. Ambrosius. Remote identification of polar bear maternal den habitat in northern Alaska. Arctic 2001; 54:115–121.

8. Durner, G.M., S. C. Amstrup, and A. S. Fischbach. Habitat characteristics of polar bear terrestrial maternal den sites in northern Alaska. Arctic 2003; 56:55–62.

9. Durner, G.M., S. C. Amstrup, and K. J. Ambrosius. Polar bear maternal den habitat in the Arctic National Wildlife Refuge, Alaska. Arctic 2006; 59: 31–36.

10. Fischbach, A. S., S. C. Amstrup, and D. C. Douglas. Landward and eastward shift of Alaskan polar bear denning associated with recent sea ice changes. Polar Bio. 2007; 30:1395–1405.

11. Hansson, R., and Thomassen, J. Behavior of polar bears with cubs in the denning area. Int. Conf. on Bear Res. and Manage. 1983; 5:246–254.

12. Larsen, T. Polar bear denning and cub production in Svalbard, Norway. J. Wildl. Manage. 1985; 49:320–326.

13. Linnell, J.D.C., J. E. Swenson, R. Andersen, and B. Barnes. How vulnerable are denning bears to disturbance? Wildl. Soc. Bull. 2000; 28:400–413.

14. Ovsyanikov, N. Den use and social interactions of polar bears during spring in a dense denning area on Herald Island, Russia. Int. Conf. Bear Res.Manage. 1998; 10: 251–258.

15. Robinson, R., T. S. Smith, R. T. Larsen and B. J. Kirschhoffer. Factors influencing polar bear den detection using forward-looking infrared imagery. BioSci. 2014; 64: 735–742.

16. Rode, K. D., J. W. Olson, D. Eggett, D. C. Douglas, G. M. Durner, T. C. Atwood, E. V. Regher, R. S. Wilson, T. S. Smith, M. St. Martin. Den phenology and reproductive success of polar bears in a changing climate. J. Mamm. 2018; 99:16–26.

17. Serreze M. C., A. P. Barrett, and J. Stroeve. Recent changes in tropospheric water vapor over the Arctic as assessed from radiosondes and atmospheric reanalyses. J. Geophys. Res. 1998; 117, D10104, doi:10.1029/2011JD017421.

18. Smith, T. S., S. T. Partridge, S.C. Amstrup, and S. Schliebe. Post-den emergence behavior of polar bears (*Ursus maritimus*) in Northern Alaska. Arctic 2007; 60: 187–194.

19. Smith, T. S., J. A. Miller and C. S. Layton. A comparison of methods to document activity patterns of post-emergence polar bears (*Ursus maritimus*) in northern Alaska. Arctic 2013; 66: 139–146.

20. U.S. Fish and Wildlife. Polar Bear (*Ursus maritimus*) Conservation Management Plan, Final. 2016. U.S. Fish and Wildlife, Region 7, Anchorage, Alaska. 104 pp.

21. USGS Alaska Science Center, Polar Bear Research Program. Denning phenology, den substrate, and reproductive success of female polar bears (*Ursus maritimus*) in the Southern Beaufort Sea 1986-2013 and the Chukchi Sea 1987-1994: U.S. Geological Survey data release, https://doi.org/10.5066/F7DF6PC9.

22. Uspenski, S.M., and Kistchinski, A.A. New data on the winter ecology of the polar bear (*Ursus maritimus*) on Wrangel Island. International Conference on Bear Research and Management 1972; 2: 181–197.

23. York, G., S. C. Amstrup, and K. Simac. Using forward looking infrared (FLIR) imagery to detect polar bear maternal dens: operations manual. USGS Report submitted to USDOI-MMS, 2004; 58 pp.

